# Volatile anaesthetics modulate voltage-gated sodium channel function at a site critical for gating

**DOI:** 10.1101/2024.11.04.621342

**Authors:** David Hollingworth, Karl F. Herold, Geoff Kelly, Vitaliy B. Mykhaylyk, Donghang Zhang, B. A. Wallace, Hugh C. Hemmings

**Author notes:** Contributed equally to the work (David Hollingworth, Karl F. Herold).

## Abstract

Voltage-gated sodium channels (VGSCs), key mediators of excitability and synaptic transmission, are established and functionally relevant targets for volatile anaesthetic (VA) action. Using the structurally homologous prokaryotic VGSCs NavMs and NaChBac as models, we present a structure-function analysis of VGSC-VA interactions. We report that multiple VAs compete for binding sites on NavMs, and that these direct interactions mediate functional effects of sevoflurane on NavMs that mirror those attributed to VA effects in eukaryotic VGSCs, including human isoforms. Using X-ray crystallography, we determined the first atomic-resolution structure of a VA bound to a VGSC, showing sevoflurane displacing lipids to bind in an intramembranous hydrophobic pocket of NavMs. A conserved tyrosine residue within this binding site is critical for channel gating, and its substitution with alanine abolishes sevoflurane binding and selectively eliminates the characteristic anaesthetic-induced hyperpolarising shift of steady-state inactivation that reduces neuronal excitability at physiological membrane potentials. Finally, we provide evidence supporting VA action at the conserved sites in human VGSC isoforms. These findings define the first VA binding site in a VGSC. A membrane-mediated access pathway to the binding site leads to negative modulation of channel function that reduces neuronal activity and excitatory synaptic transmission in general anaesthesia.

## Introduction

Since the introduction of diethyl ether as the first general anaesthetic in the 1840s, volatile anaesthetics (VAs) have become established as essential medicines for enabling painful procedures by producing amnesia, unconsciousness and immobility. Nonflammable halogenated ethers have since replaced diethyl ether, with desflurane, isoflurane and sevoflurane, the dominant VA used in developed countries, in current use^1,2^. Despite their clinical importance, the mechanisms by which VAs produce anaesthesia remain poorly understood^3^, although the positive correlation between VA potency and lipophilicity has implicated hydrophobic binding processes^4^.

The principal neuronal targets of VAs are membrane proteins, particularly ligand-gated and voltage-gated ion channels^3-6^. Their relatively low-affinity interactions with relevant target proteins modulate the neural networks responsible for consciousness, memory, and nocifensive responses through suppression of excitatory and enhancement of inhibitory synaptic transmission^7^. As these membrane proteins are essential for the physiological functions of excitable cells in the nervous system, skeletal muscle, and cardiovascular systems, such interactions are also implicated in the serious adverse side effects associated with VA use, including neurotoxicity, cognitive dysfunction, respiratory depression, and cardiovascular complications^8,9^. Identifying and characterising functionally important anaesthetic binding sites in these target molecules is crucial for understanding their effects to help define their molecular pharmacology, thus enabling the design of more selective VAs with fewer side effects.

Voltage-gated sodium channels (VGSCs) are critical to cellular excitability, and are important targets for the neurophysiological effects of VAs^3,10-16^. The nine human VGSC isoforms (Nav1.1 - Nav1.9) are expressed primarily in excitable tissues^17^, where they drive action potentials and shape excitability by cycling through resting (closed, non-conducting), open (conducting) and inactivated (closed, non-conducting) conformational states in a voltage-dependent manner^17^. Sevoflurane, desflurane, isoflurane and halothane all affect neuronal VGSCs similarly at clinically relevant concentrations by inhibiting peak Na^+^ current (*I*_Na_), accelerating current decay, shifting steady-state inactivation to more negative membrane potentials and delaying recovery from inactivation^13-16,18-20^. Preferential interactions with the inactivated state^11,16,18,21,22^ contribute to reduced cellular excitability,^21,23,24^ exocytosis^25,26^ and neurotransmitter release^25,27^, both neurophysiological correlates of the anaesthetic state.

While the functional effects of VAs on VGSCs are relatively well established, understanding the molecular mechanisms underlying these effects has been limited by the lack of atomic-resolution structural data to define their binding sites. Obtaining high-resolution structural data on eukaryotic VGSCs is challenging as these structurally complex membrane proteins are refractory to X-ray crystallography. While several cryo-electron microscopy (cryo-EM) structures of eukaryotic VGSCs are available, no cryo-EM structure of a VA bound to any protein has been reported. This is likely due to the inherent challenges in imaging small, volatile, and weakly binding molecules using this technique.

Bacterial homologues have been used as effective models of eukaryotic ion channels in structural studies of ion channel-drug interactions to determine the molecular basis for drug effects. For example, X-ray crystallography has been used to identify and characterise binding sites for the VAs isoflurane and desflurane in bacterial homologues of pentameric ligand-gated ion channels including GABA_A_ receptors, another major target for VAs^28,29^. Bacterial VGSCs (BacNavs) share similar structural and functional properties with their eukaryotic counterparts^30,31^, and the BacNav NaChBac, from *Bacillus halodurans*, is inhibited by VAs at clinical concentrations^13,32,33^. Binding sites for VAs in NaChBac have been predicted using molecular dynamics simulations and nuclear magnetic resonance (NMR) spectroscopy^32,34,35^, but NaChBac has not been crystallised, so no atomic resolution structures exist to substantiate these putative VA binding sites. In contrast, NavMs, from *Magnetococcus marinus*, crystallises to high resolution and has provided an excellent BacNav model for understanding drug actions on human VGSC^36-38^, making it a viable candidate for this study.

NavMs has the typical BacNav structure of four identical ∼30 kDa monomeric subunits that assemble to form a homotetrameric channel^39^. Each subunit consists of 6 transmembrane helices (S1-S6, labelled 1-6 in Fig. 1a) that form channels with four peripheral voltage-sensing domains (VSDs; helices S1-S4), each connected by a linker helix (the S4-S5 linker helix) to a sodium ion (Na^+^) conducting pore module (PM, consisting of helices S5-S6 and their interconnecting loops from all four subunits) in a domain-swapped arrangement where the VSD from one subunit packs against the PM from the adjacent subunit^31^ (Fig. 1b).

**Fig. 1.**
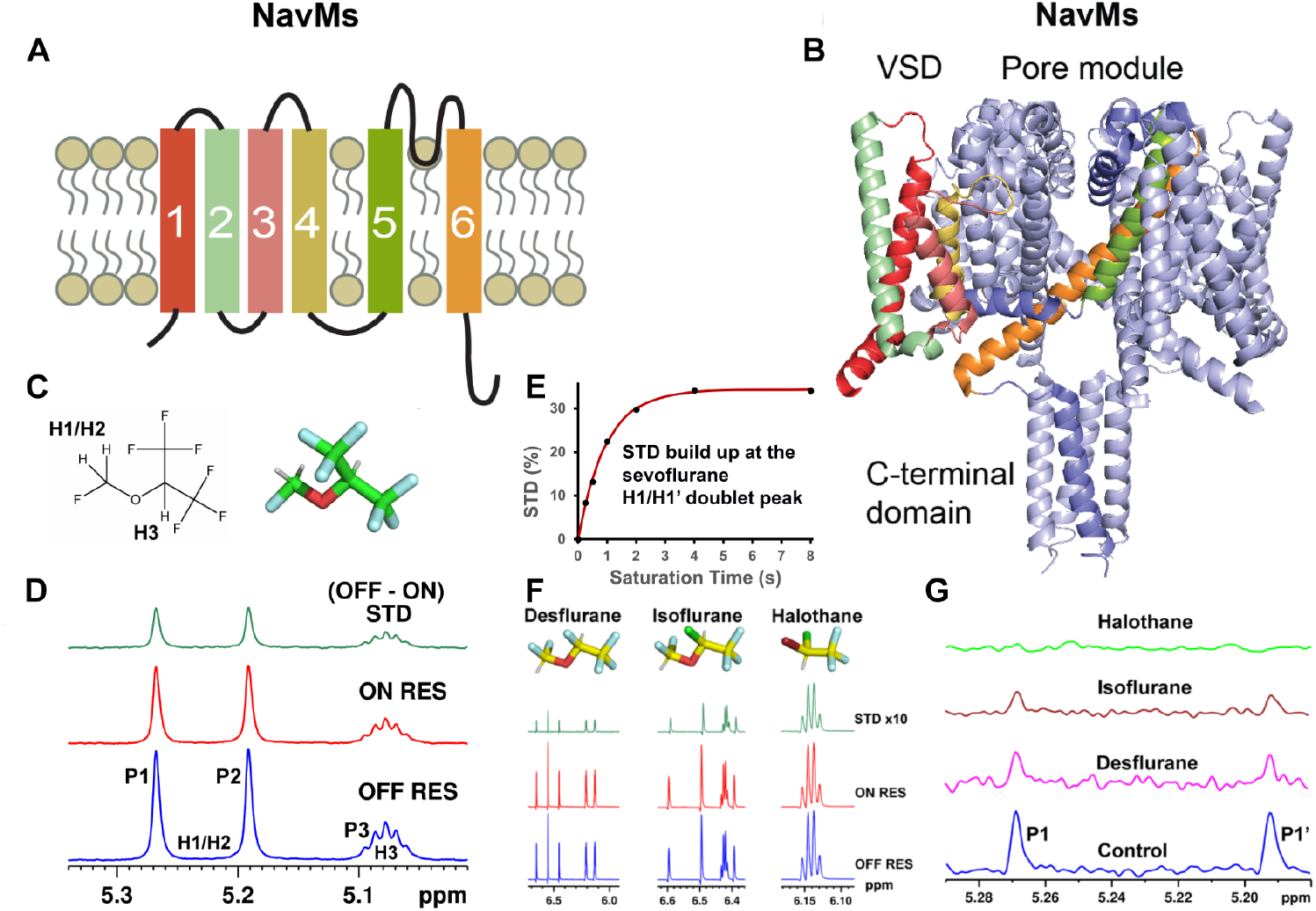
The homotetramer NavMs interacts directly with multiple volatile anaesthetics. **a**, Membrane topology of NavMs in which 6 transmembrane helices (1-6) make up the monomeric subunit. **b**, NavMs tetramers form the functional domain-swapped channel with one subunit having coloured helices as in (**a**). **c**, Sevoflurane chemical structure with labelling of the three sevoflurane protons as used in this study *(left)* and in stick representation with green backbone as used in this study *(right)*. **d**, Stacked NMR spectra show STD *(top*, green) produced after 4 s saturation at the OFF (*bottom*, blue) and ON (*middle*, red) resonances, showing that Saturation Transfer Difference (STD) is produced at all sevoflurane proton peaks. P1/P1’ is a single split peak formed by splitting of the peak for the equivalent H1 and H1’ protons shown in (**c**) to give a doublet peak due ^2^J_HF_ coupling with the fluorine of the CH_2_F-group, and P2 is produced by the H3 proton shown in (**c**) as a broad septet peak due to ^3^J_HF_ coupling with the six fluorine atoms of the two neighbouring CF_3_-groups. (**e**) Saturation time-dependent build-up of STD at the P1/P1’ resonances continues until plateau (STDmax) at ∼ 4 s. STD(%) is calculated as (I_off – I_on) / I_off × 100, where I_off and I_on are the integrals of the sevoflurane proton peaks in the OFF RES (irradiation −20 ppm) and the ON RES (−0.5 ppm) spectra at the indicated saturation times. Plot shows STD *versus* time at P1/P1’ with the experimental values fitted to Equation 1 (see *Methods*). **f**, Similar stacked NMR STD spectra as shown in (**d**) for desflurane (*left*), isoflurane (*middle*) and halothane (*right*) show STD for all three VAs indicating direct interaction (a stick drawing of each VA is provided at the top of each panel with yellow backbone for clarity). **g**, VAs compete for binding sites on NavMs. Stacked spectra shown at the sevoflurane H1/H1’ doublet peak at STDmax show that the STD produced without competitor VA (*bottom* spectrum) is reduced by saturating concentrations of desflurane (pink) and isoflurane (brown), and abolished by halothane (light green).

Outward movement of the S4 helix within the VSD is transmitted via the S4–S5 linker to the pore module (PM), inducing conformational changes that drive the transition between closed and open channel states. Lateral fenestrations in the PM form intramembrane openings to the channel pore, providing a membrane accessible pathway for drugs that can block channels from their closed resting state^40^. While BacNavs share the fundamental tetrameric architecture of all VGSCs, eukaryotic channels consist of a single amino acid chain (∼210 kDa) with four non-identical domains (DI-DIV) replacing the four identical BacNav subunits to create a pseudo-tetrameric structure.

We present the first atomic-resolution crystal structure of the VA sevoflurane bound to the voltage-gated sodium channel NavMs, revealing a membrane-accessible hydrophobic pocket as a selective binding site. We further demonstrate that sevoflurane, along with desflurane, isoflurane, and halothane, compete for common binding sites on VGSCs to produce their functional effects. Binding of sevoflurane at this membrane-accessible site selectively stabilises the inactivated channel state, producing a characteristic hyperpolarising shift in steady-state inactivation, while additional, unidentified sites appear to mediate other VA-induced effects observed in wild-type channels. Substitution of a conserved tyrosine residue in this pocket with alanine disrupts the sevoflurane interaction at this site and abolishes the shift in steady-state inactivation, highlighting its critical role. To bridge these findings to homologous sites in human VGSCs, we show that sevoflurane’s functional effects on NavMs are preserved in an NavMs mutant with the predicted human binding site residues, and *in silico* docking to neuronal human Nav1.1 support binding within the corresponding hydrophobic pockets. VAs stabilise the inactivated state of human VGSCs, a central mechanism in depressing excitatory neurotransmission, a hallmark of clinical anaesthesia^20,41^, which supports the physiological importance of this binding site *in vivo*.

## Results

### Sevoflurane interacts directly with NavMs

We investigated direct interaction between VAs and NavMs using saturation transfer difference nuclear magnetic resonance (^1^H STD NMR) spectroscopy^42^ to identify weak protein-ligand interactions such as those reported for VA binding to ion channel targets, including NaChBac^34^. These interactions are reported through the reduction in the intensities of ligand ^1^H NMR signals, caused by internuclear saturation transfer from selectively irradiated protein to ligand when in close (distances < 7 Å) and repetitive contact^43^. We probed for interactions between NavMs and sevoflurane, which contains 3 protons (Fig. 1c), using a clinically relevant aqueous sevoflurane concentration of 0.57 mM (equivalent to 2 MAC, twice the 50% effective dose or minimum alveolar concentration [MAC]); this concentration is used in the subsequent functional experiments throughout this study).

Selective irradiation of the NavMs ^1^H signal at −0.5 ppm (ON resonance) resulted in saturation transfer from NavMs to sevoflurane reducing the intensities of all three sevoflurane ^1^H spectral signals (Fig. 1d, *middle*); this did not occur when irradiation was performed at a control resonance outside the NavMs ^1^H spectrum (−20 ppm, OFF resonance, Fig. 1d, *bottom*). The difference (OFF resonance – ON resonance) resulted in the saturation transfer difference spectrum (STD, Fig. 1d, *top*), which increased in a time-dependent manner until STDmax (∼33%) was achieved at plateau (Fig. 1e). No STD occurred when the experiment was conducted in the absence of NavMs (Supplementary Information, Fig. S1). Together, these data identify that direct interaction occurs between NavMs and sevoflurane.

To investigate whether the modern VAs desflurane, isoflurane, and halothane directly interact with NavMs, and whether they occupy the same sites as sevoflurane despite structural differences, we performed competition STD experiments by introducing buffer-saturating concentrations of a competitor VA alongside 1 μM NavMs and 2 MAC sevoflurane. Saturating concentrations of VAs were used to observe their effects on the sevoflurane STD signals, considering that this technique relies on fast-exchanging ligands. This led to ratios of competing VA to sevoflurane of ∼35:1 for desflurane, ∼40:1 for isoflurane and ∼80:1 for halothane. STD was produced at the proton resonances of each of the competitor VAs (shown at STDmax in Fig. 1f), indicating their direct interaction with NavMs. Under these experimental conditions, but in the absence of any competitor VA, sevoflurane produced time-dependent STD up to a plateau (STDmax of ∼8% after 8 sec saturation time, control, shown at the P1/P1’ resonances; Fig. 1g, blue). The presence of desflurane caused ∼50% reduction in sevoflurane STD at STDmax (Fig. 1g, pink), isoflurane ∼71% reduction (Fig. 1g, brown), with halothane completely abolishing sevoflurane STD (Fig. 1g, light green), demonstrating that all three VAs compete with sevoflurane for binding sites on NavMs. The observation that halothane, a chemically distinct VA, fully displaces sevoflurane from NavMs binding suggests that all four VAs target shared sites, though desflurane and isoflurane would require higher, experimentally unattainable concentrations to completely eliminate sevoflurane’s STD signal.

### Sevoflurane has similar effects on bacterial and mammalian VGSC function

VAs affect mammalian, including human, VGSCs at clinically relevant concentrations by inhibition of peak Na^+^ current (*I*_Na_), accelerated current decay, a hyperpolarising shift in the voltage-dependence of steady-state inactivation and slowed channel recovery from inactivation^13,18^, with no effects on activation. We tested the effects of sevoflurane on NavMs function using whole-cell patch clamp electrophysiology in human embryonic kidney 293T (HEK293T) cells transiently transfected to express NavMs. Sevoflurane reduced peak *I*_Na_ from the resting state (Fig. 2a,b,d), accelerated current decay (Fig. 2c), shifted the voltage-dependence of steady-state inactivation toward more hyperpolarised potentials without altering activation kinetics (Fig. 2d), and slowed recovery from inactivation (Fig. 2e,f, Supplementary Information Table S1). These functional effects of VAs mirror those on human VGSC subtypes, and are consistent with multisite VA binding to VGSCs observed in molecular dynamics studies^32,34,35^.

**Fig. 2.**
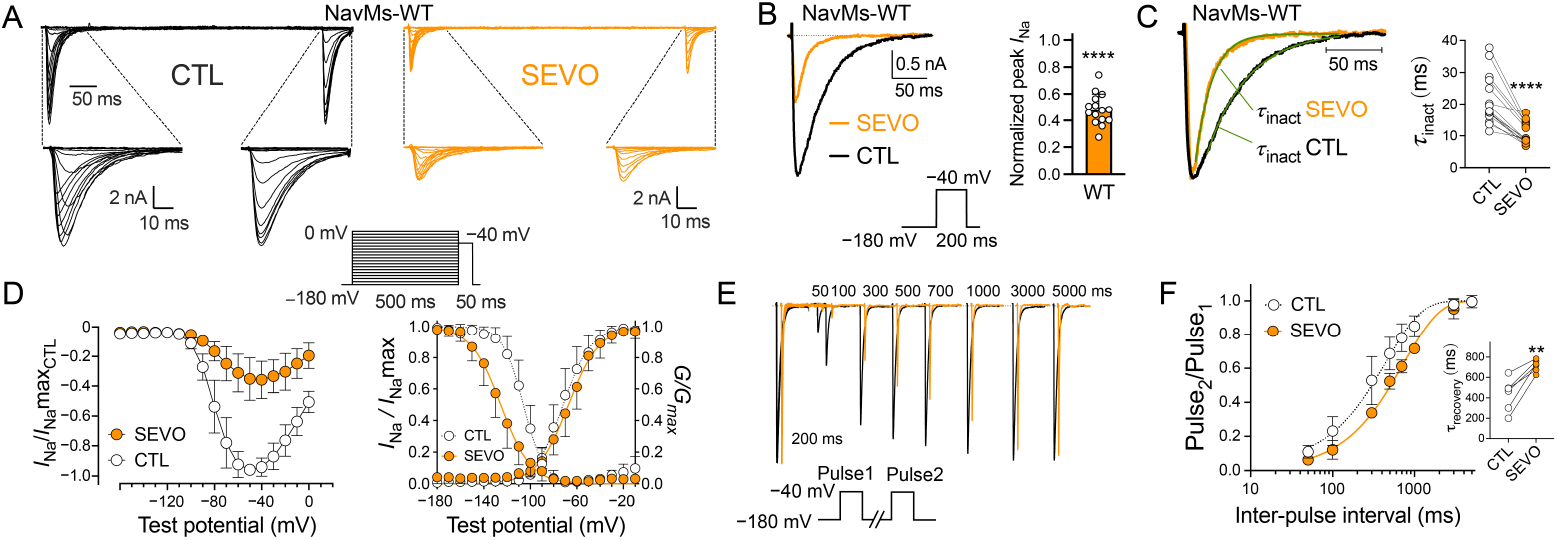
Sevoflurane modulates NavMs function. **a**, Representative families of whole-cell inward Na^+^ current (*I*_Na_) evoked using the stimulation protocol shown *(inset*) from a holding potential *(V*_h_) of −180 mV in the absence (CTL; *left*, black traces) or presence of 0.57 mM (2 MAC) sevoflurane (SEVO; *right*, orange traces). **b**, Sevoflurane inhibition of NavMs peak *I*_Na_. Representative traces of peak *I*_Na_ evoked by a single pulse (*inset*) from a holding potential (*V*_h_) of −180 mV in the absence (CTL; black trace) or presence of sevoflurane (SEVO; orange trace). Normalised peak *I*_Na_ was reduced to 0.48±0.12 by 0.57 mM (2 MAC) sevoflurane (****P<0.0001, n=15). **c**, Sevoflurane reduced the time constant of current decay (τ_inact_). Traces in (**b**) were normalised and data for current decay were fitted to a mono-exponential equation (green curves) to calculate τ_inact_. Sevoflurane accelerated current decay: τ_inact_ was reduced by ∼2-fold from 21.3±7.9 ms (CTL; white circles, *right* panel) to 10.6±3.4 ms (SEVO; orange circles, ****P<0.0001, n=15). **d**, Current-voltage relationship of channel activation (I-V curve, *left* panel), normalised conductance (*G*/*G*_max_) and inactivation (*I*_Na_/*I*_Na_max; *right* panel). Sevoflurane did not significantly affect the voltage dependence of activation (*V*_½act_ −71.5±7.6 mV for CTL [white circles] *vs*. −66.7±6.6 mV for SEVO [orange circles], P=0.0695, n=8). Sevoflurane shifted the voltage dependence of half-maximal inactivation (*V*_½inact_) towards more negative potentials by −19.4±6.0 mV (−106±6.4 mV for CTL [white circles] *vs*. −126±8.2 mV for SEVO [orange circles], ****P<0.0001, n=8). **e** and **f**, Sevoflurane slowed Na^+^ channel recovery from inactivation. A two-pulse protocol was used from a holding potential (*V*_h_) of −180 mV with two 200 ms test pulses to −40 mV separated by an interpulse interval of 50 to 5000 ms (*inset*). Peak *I*_Na_ of the second test pulse was normalised to the first (Pulse_2_/Pulse_1_) and plotted against the interpulse interval. Data were fitted to a mono-exponential equation to calculate the time constant τ. **e**, Overlaid normalised macroscopic Na^+^ current traces of two pulses with increasing interpulse intervals in the absence (CTL; black traces) or presence (SEVO; orange traces) of 0.57 mM (2 MAC) sevoflurane. The sevoflurane traces were plotted with an *x*-axis offset of +30 ms to allow visual comparison between the current dynamics. Peak *I*_Na_ of the second test pulse recovers more slowly with increasing interpulse intervals in the presence of sevoflurane. **f**, Fitted data in the absence (CTL; white circles) or presence (SEVO; orange circles) of sevoflurane. *Inset* shows recovery time constants τ derived from the fitted data. Time constant τ increased from 432±158 ms to 714±59 ms in the presence of sevoflurane, slowing recovery from inactivation (**P=0.0023, n=6). Data shown as mean±SD; drug effects *vs*. control were tested by paired two-tailed Student *t*-test (*P<0.05; **P<0.01; ****P<0.0001).

### Identification of a sevoflurane binding site in NavMs at atomic resolution

No atomic-resolution structure of a VA bound to any voltage-gated ion channel has been reported to date, so the binding site(s) underlying VA-VGSC interactions inferred from NMR, MD simulations, and electrophysiological studies have not been identified. We utilised NavMs F208L, which is functionally indistinguishable from WT NavMs but crystallises more reliably and has been used previously in structure-function studies of VGSC-drug interactions^36-38^. Apo-NavMs F208L crystals were grown and incubated with sevoflurane-saturated mother liquor, then harvested after increasing incubation times (denoted as NavMs-SEVO crystals). X-ray diffraction data were collected for both NavMs-SEVO crystals and apo-NavMs crystals (apo-NavMs). Prolonged incubation periods with sevoflurane led to diminished or no X-ray crystal diffraction, but shorter incubation times (<20 min) resulted in some crystals that diffracted to the high resolution of apo-NavMs crystals. Comparison of multiple datasets from control and sevoflurane-exposed crystals showed that while the protein structures were identical (rmsd <0.3 Å), a marked change in the shape of non-protein electron density occurred in a hydrophobic pocket below the side chain of residue Y143, located on the S5 helix of the channel pore domain. Polder-OMIT maps^44^, generated to isolate these ligand electron densities minimised for interference from bulk solvent effects, showed that apo-NavMs crystals had a short, elongated stretch of density attributed to part of the alkyl chain of lipid or detergent (Fig. 3a, *left*), while the electron density at this location in NavMs-SEVO crystals was ellipsoidal (Fig. 3a, *middle*) with a shape into which a single sevoflurane molecule could be fit in several orientations.

**Fig. 3.**
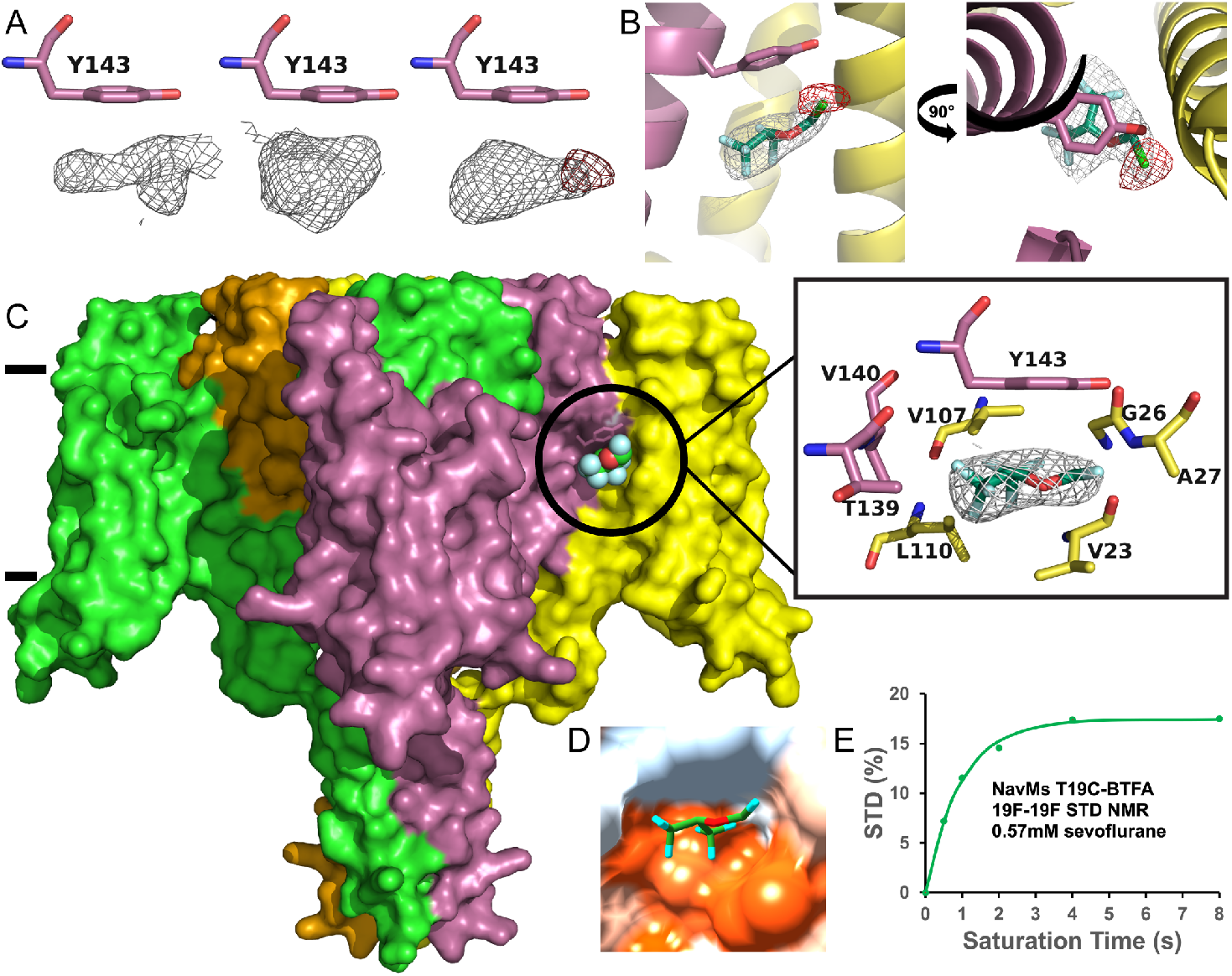
Sevoflurane binds to NavMs, displacing lipid at a membrane-exposed site. **a**, Polder OMIT map density showing the changes in the morphology of electron density (grey) below the helix S5 residue Y143 in apo-NavMs F208L crystals (*left*, contoured @2.5σ), NavMs-SEVO crystals (*middle*, contoured @4σ) and chloroSEVO crystals (*right*, contoured @4σ), also showing the anomalous signal for the chlorine atom in chloroSEVO crystals (red density, contoured @4σ). **b**, Two views of chloroSEVO (stick representation in grey electron density contoured at 1σ) in its binding site located between the S1 and S4 helices of one subunit (*yellow*) and the S5 helix from the adjacent subunit (magenta). Anomalous chlorine electron density (red, 4σ) allows for accurate positioning of chloroSEVO into the density. **c**, Space filling model (*left*, coloured by subunit) showing sevoflurane (sphere representation) in its intramembrane binding site below Y143 (membrane boundaries indicated with black lines on *left* side of image). Close-up of the sevoflurane binding site (*right inset*) showing residues (coloured by subunit) involved in binding to sevoflurane (stick) with sevoflurane electron density contoured at 1σ. **d**, Hydrophobic surface representation of sevoflurane (stick) bound in the hydrophobic intramembrane pocket created by the binding site residues (coloured by hydrophobicity *red* - highest to *blue* - lowest). **e**, ^19^F-^19^F STD build-up curve at the sevoflurane fluorine resonance (−74.7 ppm) shows the relevance of the site for sevoflurane binding at the clinically relevant concentration of 0.57 mM (2 MAC). STD (%) is calculated as (I_off – I_on) / I_off × 100, where I_off and I_on are the integrals of the sevoflurane fluorine peak at −74.7 ppm in the OFF RES spectrum (where irradiation is at 0 ppm, empty of fluorine signal) and the ON RES spectrum (where irradiation is at −83.8 ppm, which contains the unique fluorine signal from BTFA-labelled NavMs T19C) at the indicated saturation times. Curve produced from experimental values fit using Equation 1 (see *Methods*).

### Orientation and binding site interactions of sevoflurane in the binding pocket

To identify this electron density unambiguously as sevoflurane and to orientate sevoflurane into this binding site, we carried out the same crystal exposure experiments replacing sevoflurane (2-(fluoromethoxy)-1,1,1,3,3,3-hexafluoropropane) with its monochloro-substituted analogue 2-(chloromethoxy)-1,1,1,3,3,3-hexafluoropropane (chloroSEVO). This analogue retains anaesthetic properties but contains a fluorine-to-chlorine substitution on the fluoromethyl moiety of sevoflurane^45^. Chlorine is a weak anomalous scatterer that can be detected using long-wavelength X-ray crystallography^46^. Analysis of data collected on three NavMs-chloroSEVO crystals showed the same ellipsoidal density found below Y143 in the electron density map of NavMs-SEVO, with the anomalous data containing a signal within the density (Fig. 3a, *right*) that overlapped across all three crystal datasets (Supplementary Information, Fig. S2), but which was not present in the anomalous data collected from apo-NavMs crystals. These data confirmed sevoflurane binding to this site in the NavMs-SEVO crystals, with the anomalous chlorine signal allowing placement of chloroSEVO, and by inference sevoflurane, into this binding site (Fig. 3b and 3c).

Sevoflurane binds in a preformed pocket of NavMs (Fig. 3c and 3d) through relatively long-range (mostly >3.5 Å) hydrophobic interactions with V23, G26, and A27 from helix S1 and V107 and L110 from helix S4 of the VSD of one NavMs subunit, and with T139, V140 and Y143 from the S5 helix of the PM from the adjacent NavMs subunit (Fig. 3c, boxed). One trifluoromethyl group of sevoflurane interacts with G26 and the side chains of V23, V107, and L110, and the other trifluoromethyl group interacts with the side chains of T139 and V140. The fluorine of the fluoromethyl group contacts A26. Atoms from all three groups make multiple contacts with Y143, making it the primary binding residue within the binding site. Sevoflurane binding displaces the alkyl tail of a lipid (or hega-10 detergent, which can replace lipid during purification) to fill the natively membrane-embedded hydrophobic cavity, disrupting the protein-lipid environment, which might be required for normal channel function^47^. This pocket would be dynamic during channel gating, as the S4 helix moves in response to the membrane potential between polarised and depolarised channel states.

Consequently, the sevoflurane binding site found in the inactivated channel state of our structure would not be present in the resting (polarised) channel state.

Previous MD simulations identified multiple potential VA binding sites within VGSCs^32,35^. This binding site was not identified in these studies, but, as this is the first binding site to be determined at atomic resolution, for simplicity, we term this binding site BS1 for the remainder of this study.

The highest resolution structures of apo-NavMs and NavMs-SEVO collected, both solved up to a resolution of 2.2 Å, have been deposited in the Protein Data Bank (apo-NavMs PDB ID: 9GTQ, NavMs-SEVO PDB ID: 9GV1).

### Sevoflurane binds BS1 at a clinically relevant concentration

The crystal structure of sevoflurane bound to NavMs was determined under conditions of saturated sevoflurane concentration. To determine whether sevoflurane interacts with BS1 at the same clinically relevant concentration used in our functional studies, we employed a fluorine-modified STD NMR technique (^19^F-^19^F STD NMR) in which proteins are site-selectively fluorinated at cysteine residues introduced near predicted binding sites to report on fluorinated ligand interactions with those sites^34^. NavMs contains one native cysteine residue, which was mutated to alanine (C52A) prior to incorporating the T19C mutation to make NavMs C52A/T19C. We chose to mutate and modify T19 because the fluorinated probe introduced at this position would be too far away to report on any of the potential VGSC-VA binding sites previously proposed from MD simulations using NaChBac^34,35^, allowing selective information for sevoflurane binding to BS1 where fluorine atoms of sevoflurane are in close proximity (∼6 Å) to the probe fluorines.

NavMs C52A/T19C expression was extremely low (∼15x lower compared to WT NavMs), but the gel filtration elution profile showed a peak that matched that of WT NavMs, indicative of an equally well-folded protein. Fluorination was performed using the cysteine-specific alkylating agent 3-bromo-1,1,1-trifluoroacetone (BTFA) which conjugates a trifluoromethyl-containing group onto free cysteines. Selective irradiation at the trifluoromethyl resonance of BTFA-labelled NavMs C52A/T19C (at −83.8 ppm) incubated with 2 MAC sevoflurane produced time-dependent STD build up at the two sevoflurane fluorine resonance peaks present in the 1D ^19^F spectrum, (at −74.7 ppm representing the six-equivalent fluorine’s in the hexafluoroisopropyl group of sevoflurane (Fig. 3e), and at −152.2 ppm representing the single fluorine in its fluoromethyl group), indicating that sevoflurane interacts with BS1 at this clinical concentration. Relative STDmax produced at the ^19^F peak of the hexafluoroisopropyl group fluorine resonance was greater (17.5%, Fig. 3e) compared to that produced at the fluoromethyl group fluorine (3.5%, Supplementary Information, Fig. S3). This result is explained by the orientation of sevoflurane in BS1, where fluorine atoms of the hexafluoroisopropyl group of sevoflurane would be >3 Å nearer to the T19-fluorine probe compared to the fluoromethyl fluorine, which is located towards the end of the range of detection of this technique.

### Y143 is necessary for both sevoflurane binding to BS1 and normal channel function

From our structural studies, the Y143A mutation of NavMs would be expected to abolish sevoflurane binding at BS1, and therefore would facilitate a detailed characterisation of the contribution of sevoflurane binding at this site from the background of the overall functional effects produced by sevoflurane binding to NavMs. However, the approach of introducing Y143A into the double mutant (producing NavMs C52A T19C Y143A) and using ^19^F-^19^F STD NMR as above, to selectively explore binding at this site, was not possible as the expression level of this protein was too low for practical use. Instead, we tested for the effect of this mutation using the single mutant NavMs Y143A and ^1^H STD NMR. NavMs Y143A had good expression and purified with a peak during gel filtration at the same retention time as WT NavMs. ^1^H STD NMR of NavMs Y143A and WT NavMs with 2 MAC sevoflurane showed that NavMs Y143A produced lower STD compared to WT NavMs (STDmax, Y143A = 29.6±2.3, WT = 34.2±0.9, *P<0.034, n=3, at the P1-P1’ doublet peak in Fig. 4a). This is consistent with reduced sevoflurane binding to NavMs Y143A compared to WT NavMs caused by the mutation eliminating sevoflurane binding to BS1.

**Fig. 4.**
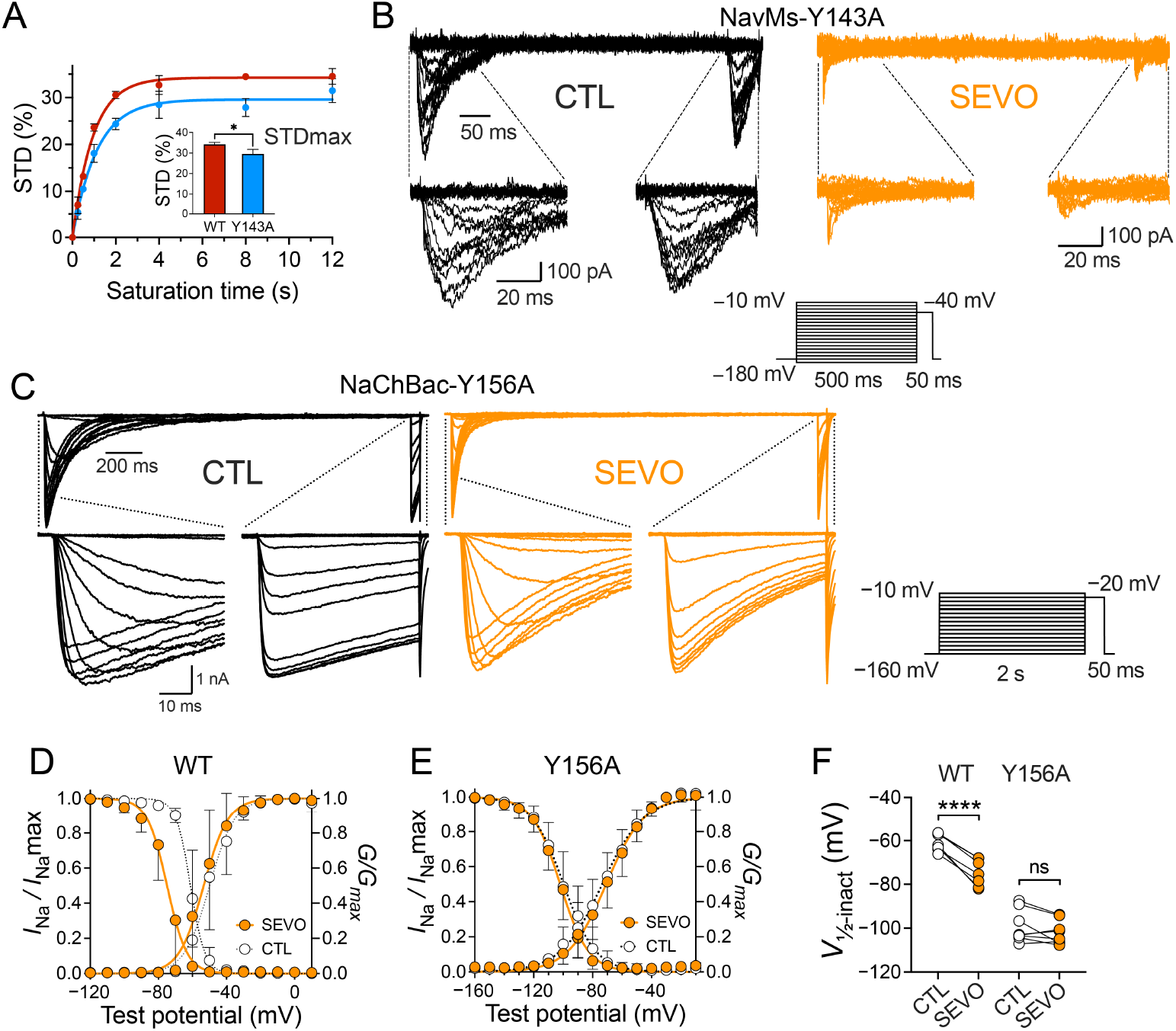
Functional effects of selectively abolishing sevoflurane binding using tyrosine to alanine mutations in NavMs (Y143A) and NaChBac (Y156A). **a**, STD build up curves produced at the proton P1/P1’ resonances by 0.57 mM (2 MAC) sevoflurane for WT NavMs (*red*) and NavMs Y143A (*blue*). Y143A had a reduced STDmax (%) compared to WT (*inset*, Y143A 29.6±2.3, WT 34.2±0.93, *P<0.05, unpaired two-tailed Student *t*-test, n=3), consistent with reduced interaction at the Y143A site. Curves produced from experimental values fit using Equation 1 (see *Methods*). **b**, Functional effects of sevoflurane on NavMs Y143A transiently expressed in HEK293T cells. Representative families of whole-cell inward Na^+^ currents (*I*_Na_) evoked from a holding potential (*V*_h_) of −180 mV in the absence (CTL; *left*, black traces) or presence of a reduced concentration of 0.28 mM sevoflurane (1 MAC) (SEVO; *right*, orange traces); the stimulation protocol is shown in the *inset*. The Y143A mutation caused a major leftward shift in channel gating, requiring a much more hyperpolarised holding potential for proper recording that exceeded the operational limits of the amplifier. This mutant also expressed poorly with small currents, which were further reduced by sevoflurane, precluding reliable determination of sevoflurane effects on of activation and inactivation. **c**, The homologous Y156A mutation in NaChBac had more favourable gating properties from a holding potential (*V*_h_) of −160 mV, shown in the absence (CTL; *left*, black traces) or presence (SEVO; *right*, orange traces) of 0.57 mM (2 MAC) sevoflurane; the stimulation protocol is shown in the *inset*. **d**,**e**, Comparison of voltage-dependence of activation and inactivation for WT NaChBac (**d**) and NaChBac Y156A (**e**) in the absence (white symbols) or presence (orange symbols) of sevoflurane. Current-voltage relationship of channel activation (normalised conductance; *G*/*G*_max_) and inactivation (*I*_Na_/*I*_Na_max). Sevoflurane significantly left-shifted the voltage-dependence of half-maximal inactivation (*V*_½inact_) by −13.9±3.7 mV for NaChBac WT (**d**,**f**; Supplementary Information, Table S1). This hyperpolarising shift in *V*_½inact_ was not seen for the Y156A mutant (**e**,**f**; Supplementary Information, Table S1). Data shown as mean±SD; drug effects *vs*. control were tested by paired two-tailed Student *t*-test; (*P<0.05; ****P<0.0001).

We examined the functional impact of this mutation using whole-cell patch clamp electrophysiology on HEK293T cells expressing NavMs Y143A. The Y143A mutation led to a channel that only produced small *I*_Na_ compared to WT NavMs (Fig. 4b, *left*), These currents were almost fully inhibited by sevoflurane (Fig. 4b, *right*), such that experiments necessary to characterise the consequences of sevoflurane binding to BS1 on channel function were not possible.

### Functional significance of sevoflurane-BS1 interaction revealed using NaChBac

The Y143A mutation in NavMs produced notable alterations in baseline gating compared to wild-type NavMs (Supplementary Information, Table S1), notably a marked hyperpolarising shift in the voltage-dependence of inactivation. This further lowered the already extreme negative holding voltage needed to return channels to the resting state, surpassing the technical capabilities of our amplifier and resulting in a significant decrease in *I*_Na_. NaChBac, like all BacNavs that have been identified as functional channels, has a conserved tyrosine residue (Y156) at the equivalent position to Y143 in NavMs, and has been previously used in functional studies of VA effects on VGSCs^32,33^. NaChBac displays more favourable gating kinetics than NavMs, requiring less extreme hyperpolarization to reach the resting state, with membrane potentials that more closely resemble those of mammalian VGSCs^32,33,48^. WT NaChBac produced large currents with less negative inactivation gating compared to NavMs (Fig. 4d; Supplementary Information, Table S1), as expected. Sevoflurane (2 MAC; 0.57 mM) shifted the voltage dependence of steady-state inactivation towards more hyperpolarised potentials, inhibited peak *I*_Na_, and accelerated current decay. NaChBac Y156A yielded robust currents, and affected gating by shifting both steady-state activation and inactivation curves negatively and with broader slopes compared to WT NaChBac (Fig. 4e; Supplementary Information, Table S1). Most of the effects produced by sevoflurane on WT NaChBac were maintained on NaChBac Y156A, except that the characteristic VA-induced negative shift in steady-state inactivation seen in WT channels was selectively eliminated (Fig. 4e and 4f). This property of VGSC modulation has been proposed to underlie the reduced neuronal excitability and neurotransmitter release produced by VAs^41^.

### A homologous anaesthetic binding site is conserved in human Na^+^ channels

All human VGSC isoforms contain a conserved phenylalanine residue at equivalent positions to Y143 in BS1 in the S5 helices of domains II and IV (e.g. F902 and F1682 in Nav1.1, Supplementary information, Fig. S5). Examination of the cryo-EM structure of human Nav1.1 (PDB ID: 7DTD^49^, inactivated state) identifying the presence of these residues in conserved hydrophobic pockets. To study the consequences of phenylalanine substitution at this position for channel function and sevoflurane binding, we mutated Y143 to phenylalanine in NavMs F208L. NavMs F208L/Y143F expressed similarly to NavMs F208L and purified identically. It produced crystals under the same conditions as NavMs F208L that diffracted to high resolution producing electron density maps and structures identical to those of NavMs F208L, except for the lack of density associated with the Y143 hydroxyl group. X-ray diffraction data obtained from crystals after co-crystallisation of NavMs F208L/Y143F with sevoflurane produced electron density maps that showed the same ellipsoidal density below F143 as seen below Y143 in NavMs-SEVO crystals, indicating sevoflurane binding (Fig. 5a). Whole cell patch-clamp electrophysiology on HEK293T cells expressing NavMs Y143F showed that this channel behaved like WT NavMs (Fig. 5b-e) and that sevoflurane produced similar effects to WT NavMs (Fig. 5b-e).

**Fig. 5.**
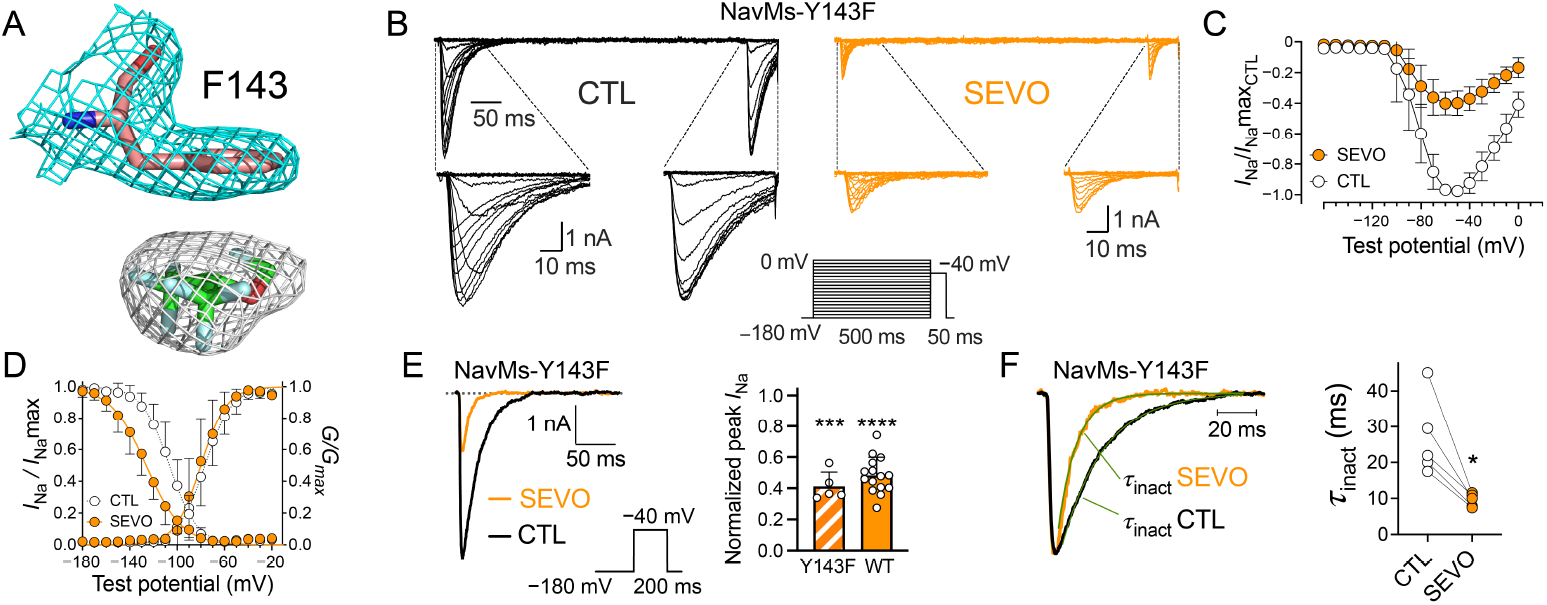
NavMs Y143F is similar to WT NavMs in both function and sensitivity to sevoflurane. **a**, Co-crystals of NavMs Y143F with sevoflurane show identical sevoflurane density to that found in NavMs-SEVO crystals. **b**, Representative families of whole-cell inward Na^+^ currents (*I*_Na_) in HEK293T cells expressing NavMs Y143F (*inset* shows stimulation protocol) from a holding potential (*V*_h_) of −180 mV in the absence (CTL; black traces) or presence of 0.57 mM (2 MAC) sevoflurane (SEVO; orange traces). **c**,**d**, Current-voltage relationships of channel activation (*I*-*V* curve; **c**), normalised conductance (*G*/*G*_max_) and inactivation (*I*_Na_/*I*_Na_max; **d**). Sevoflurane did not affect the voltage-dependence of activation (−77±9 mV for CTL; white circles, *G*/*G*_max_; *vs* −80±9 mV for SEVO; orange circles, P=0.544, n=5). Sevoflurane shifted the voltage-dependence of half-maximal inactivation (*V*_½inact_) by −20±11 mV (−107±8 mV for CTL; white circles, *I*_Na_/*I*_Na_max; *vs*. −126±8 mV for SEVO; orange circles, *P=0.0305, n=5). **e**, Sevoflurane inhibits NavMs Y143F peak *I*_Na_. Representative overlaid traces of *I*_Na_ evoked by a single pulse (*inset* shows stimulation protocol) from *V*_h_ −180 mV in the absence (CTL; black trace, *left* panel) or presence of 0.57 mM (2 MAC) sevoflurane (SEVO; orange trace). Normalised peak *I*_Na_ was reduced to 0.41±0.09 in the presence of 0.57 mM (2 MAC) sevoflurane (*right* panel; ***P=0.00014, n=5). WT NavMs data from Fig. 2b included for comparison (*right* panel). **e**, Sevoflurane increases the time course of current decay (τ_inact_) of channel inactivation in NavMs Y143F comparable to WT NavMs. The traces in (**e**) were normalised and data for current decay were fitted to a mono-exponential equation (green curves) to calculate τ_inact_. Sevoflurane accelerated current inactivation τ_inact_ from 26.8±11.2 ms (CTL; white circles; *right* panel) to 9.6±1.8 ms (SEVO; orange circles, *P=0.0172, n=5). Data shown as mean±SD; drug effects *vs*. control were tested by paired two-tailed Student *t*-test; (*P<0.05; ***P<0.001, ****P<0.0001).

### Sevoflurane binding to homologous sites in human Nav1.1

Because Y143F had no effect on sevoflurane binding to BS1 we investigated whether the phenylalanine-containing pockets in human Nav1.1 also had the potential for sevoflurane binding using *in silico* molecular docking simulations. As a control, we initially docked sevoflurane into the NavMs BS1 site. The top binding pose for sevoflurane in this simulation closely correlated with that of the crystal structure pose (Fig. 6a), binding with an estimated binding energy of −4.5 kcal/mol, indicative of the characteristic weak VGSC-VA interaction. In docking experiments at the two predicted binding pockets of human Nav1.1 (DI-DII containing F902 and DIII-DIV containing F1682), sevoflurane successfully docked into both pockets with the top hits for each simulation (Fig. 6b and 6c) showing binding poses that overlapped with sevoflurane in the NavMs-SEVO structure (Supplementary Information, Fig. S6). These simulations also predicted sevoflurane binding with characteristic weak affinities (top binding energy poses; DI-DII pocket -4.1 kcal/mol, DIII-DIV pocket -5.3 kcal/mol).

**Fig. 6.**
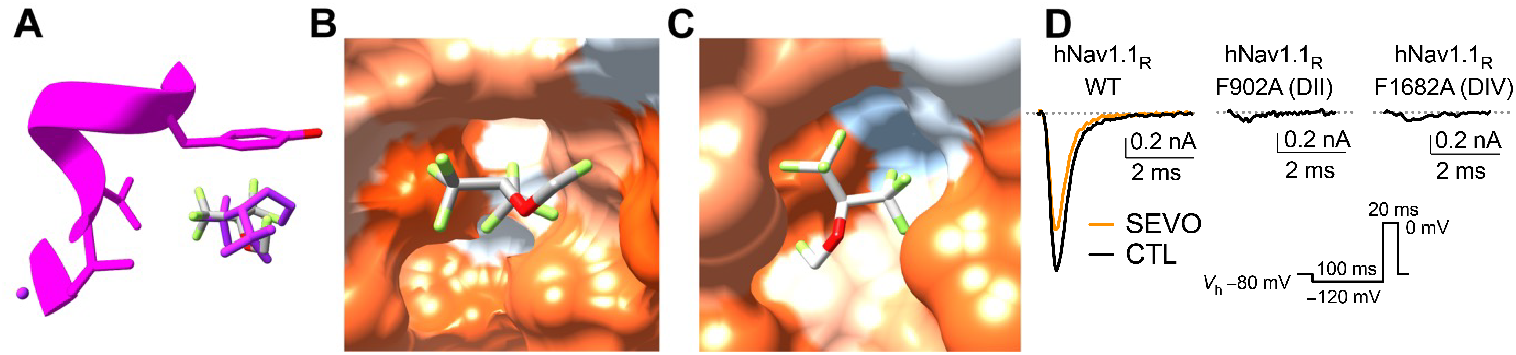
Sevoflurane interaction with equivalent sites in human Nav1.1. **a**, Molecular docking simulation of sevoflurane interaction with NavMs near the Y143 binding pocket produces top poses that overlay closely with that of sevoflurane bound in the X-ray crystal structure. Image shows the top binding pose of sevoflurane from the simulation (stick, coloured by heteroatom) overlapping closely with sevoflurane bound in the NavMs-SEVO co-crystals (*magenta*). **b**, Top pose from molecular docking simulation showing that sevoflurane (stick, coloured by heteroatom) can bind in the DI-DII pocket of human Nav1.1 (spaced filled model). **c**, Top pose from molecular docking simulation showing that sevoflurane (stick, coloured by heteroatom) can bind in the DIII-DIV pocket of human Nav1.1 (space filled model). **d**, Expression of the tetrodotoxin-resistant Nav1.1_R_ in mammalian neuronal ND7/23 cells produced a tetrodotoxin-resistant Na^+^ current (black trace, *left* panel) that was inhibited by 0.57 mM (2 MAC) sevoflurane (orange trace, *left* panel). F902A (*middle*) and F1682A (*right*) mutations introduced into human Nav1.1_R_ abolished channel function. *Inset* shows the stimulation protocol.

### F902 and F1682 in human Nav1.1 are essential for channel function

We studied the functional effects of the F902A and F1682A mutations in tetrodotoxin (TTX)-resistant human Nav1.1 expressed in rodent ND7/23 neuroblastoma cells, which are considered optimal for ion channel studies^50,51^. Modification of Nav1.1 to be resistant to TTX^52^ (F383S, referred to as Nav1.1_R_) allowed recordings of human Nav1.1 currents in isolation of the endogenous rodent Na^+^ currents that are completely blocked by 250 nM TTX. These human Nav1.1_R_-expressing cells produced strong TTX-resistant Na^+^ currents that were inhibited by sevoflurane (Fig. 6d, *left*). However, expression of the F902A and F1682A channels produced no TTX-resistant Na^+^ current, indicating that each mutation abolished human Nav1.1 function (Fig. 6d, *middle* and *right*). In contrast, mutation of S259 (S259A) in DI and L1359 (L1359A) in DIII, residues at equivalent positions to the phenylalanine residues in domains DII and DIV of human Nav1.1 (Supplemental Information, Fig. S5), had minimal effect on TTX-resistant Na^+^ currents compared to WT Nav1.1_R_ (unpublished results).

## Discussion

We identified and characterized the first atomic resolution structure of the VA sevoflurane bound to a VGSC. The VA binding site (BS1) contains a tyrosine residue critical for normal channel gating. Removal of sevoflurane binding at this site selectively abolishes the hyperpolarising shift in the voltage dependence of steady-state inactivation, a characteristic electrophysiological signature of VA effects on all VGSC subtypes, including the neuronal subtypes Nav1.1, Nav1.2, and Nav1.6^13,20^. As physiological membrane potentials in neurones maintain VGSCs in a dynamic equilibrium of their functional states, negative shifts in steady-state inactivation reduce the fraction of resting channels available for activation by increasing the number of channels in the inactivated state, which reduces neuronal excitability and stimulation-evoked synaptic vesicle exocytosis, which are neuropharmacological properties characteristic of VAs^18,41,53,54^.

The interaction between sevoflurane and NavMs at BS1 involves binding at a preformed membrane-embedded hydrophobic pocket on the lipid exposed surface of the ion channel. VAs are highly lipophilic and partition (>99%) from the aqueous phase to the membrane phase in simulations of ion channels in model membranes^32,35,55^, which provides direct access to intramembrane protein surfaces^35^ and intramembrane pockets^34,36,55^. In simulations involving VGSCs, this membrane partitioning facilitates movement of VA molecules to the membrane-embedded fenestrations through which they can move into and block the channel pore^32,35^. VA interactions with NavMs at BS1 would require a similar membrane access pathway. As the functional consequence of binding can be directly linked to inhibition of processes related to anaesthesia, this property of VA binding to VGSCs can help explain the positive correlation between VA potency and lipophilicity (the Meyer-Overton correlation). This relationship is consistent with our co-crystal structure that shows that sevoflurane must displace a lipid molecule in order to bind. Since lipids also have their own modulatory effects on VGSCs^56^, we cannot rule out that VA binding at BS1 has an indirect effect caused by displacement of lipid necessary for normal channel function. This later mechanism might explain the structural heterogeneity of the compounds which can achieve the same functional effects on VGSCs by interaction at a common site, which in our study ranged from the 2-carbon alkane halothane to the 4-carbon ether sevoflurane.

Our combined structure-function approach provides a paradigm to match specific binding events to the functional effects produced by VA binding to VGSCs with the goal of elucidating the basic molecular interactions that contribute to anaesthetic actions.

Identification of these sites is critical to further improvements in the design and development of novel anaesthetics using high throughput screening and structure-activity studies of VGSC anaesthetic binding sites^57,58^. Identification and characterisation of function-modulating anaesthetic binding sites in VGSCs could also be leveraged for drug development against VGSC channelopathies including epilepsies, arrhythmias, myotonias and neuropathic pain, where modulatory sites affecting single channel properties are considered desirable targets ^59,60^.

Eukaryotic VGSCs are essential to physiological processes that require electrical signalling.. Given their sensitivity to VAs at clinical concentrations, it is highly likely that these ion channels are affected by VAs during general anaesthesia, as supported by pharmacological studies^57,58^. Stabilisation of the inactivated state by VA binding to human VGSC at equivalent binding sites to BS1 found in BacNavs provides a structure-base mechaism for the reduction in neuronal excitability that underlies general anaesthesia^41^.

## Data availability

The atomic coordinates and crystallographic structure factors generated for Apo-NavMs and NavMs complexed with sevoflurane (NavMs-SEVO) have been deposited in the protein data bank (PDB) under the accession codes 9GTQ for Apo-NavMs [PDB DOI: https://doi.org/10.2210/pdb9gtq/pdb] and 9GV1 for NavMs-SEVO [PDB DOI: https://doi.org/10.2210/pdb9gv1/pdb] with data collection and refinement statistics details provided in Supplementary information Table S2. The PDB code of the previously published structures used in this study are for NavMs 6SX5 [https://doi.org/10.2210/pdb6sx5/pdb] and for Nav1.1 7DTD [https://doi.org/10.2210/pdb7dtd/pdb]. Source data for NMR and electrophysiology are provided as a separate data file.

## Acknowledgements

We thank the staff of Diamond Light Source (data obtained under proposal MX23853), the European Synchrotron Radiation Facility (Grenoble, France; data obtained under proposals MX2415 and MX2606) and EMBL (P13 DESY, Hamburg, Germany; data obtained under proposal MX842) for assistance and support on the beamlines used in this study. We thank Jiaxin Xiang for assistance with some of the cell culture. We thank Frances Thomas for aid with some of the images.

## Funding

US National Institutes of Health grant R01 GM58055 to HCH, a *British Journal of Anaesthesia* International Collaborative Grant to HCH and KFH, and a Rosetrees grant (ref: CF2\100001) to BAW. This work was also supported by the Francis Crick Institute through the provision of access for DH and BAW to the MRC Biomedical NMR Centre, which is funded by Cancer Research UK (CC1078), the UK Medical Research Council (CC1078), and the Wellcome Trust (CC1078).

## Author contributions

HCH, KFH and BAW conceived the project; DH and KFH designed the research; DH performed the molecular biology, protein expression and purification and chemical modifications for the structural studies; GK and DH performed and analysed NMR spectroscopy; DH performed and analysed X-ray crystallography; VBM and DH performed and analysed the long-wavelength X-ray crystallography; KFH performed the molecular biology, cell culture and electrophysiology using NavMs and NaChBac; DZ performed the electrophysiology using human Nav1.1; DH and KFH wrote the paper under the supervision of BAW and HCH.

## Conflicts of interest

HCH is editor-in-chief of the *British Journal of Anaesthesia*. The other authors have no conflicts to declare.

## Methods

### Molecular biology

Site-directed mutagenesis of the NavMs DNA sequence contained within the pET-15b plasmid vector was performed to create the mutants described in this study (Quikchange, Agilent), with the primers listed in Supplementary Information, Table S3. Plasmids generated were chemically transformed into NEB® 5-alpha competent *E. coli* (New England Biolabs) and mutation verified by automated DNA sequencing (Eurofins Genomics) before transformation of correct plasmids into Overexpress™ C41(DE3) Chemically Competent *E. coli* cells (Lucigen) for protein expression.

### Protein expression and purification

NavMs proteins were expressed in C41(DE3) *E. coli* cells grown in Luria Broth containing 100 μg/ml ampicillin, in shaking flasks at 37ºC. To induce NavMs protein expression, when the cells reached an optical density (OD600) of 0.8, 0.5 mM IPTG was added followed by further incubation for 3.5 h. Cell pellets were resuspended in 20 mM Tris pH 7.5, 150 mM NaCl, 1 mM MgCl_2_ and cells were broken by pressure. Cell debris was removed by centrifugation at 20,000x*g* for 30 min and the resulting supernatant was spun at 195,000x*g* for 2 h to pellet membranes. Protein was extracted from the membranes using a buffer containing 20 mM Tris, pH 7.5, 300 mM NaCl, 20 mM imidazole and 1.5% DDM (Anatrace) on a rotating shaker at 4ºC for 2 h. Solubilised proteins were loaded onto a 1 ml HisTrap HP column (Cytiva Life Sciences) which was washed, and detergent was exchanged on the column with buffer containing 20 mM Tris–HCl, pH 7.5, 300 mM NaCl, 50 mM imidazole and 0.52% HEGA-10 (Anatrace) prior to protein elution using the same buffer with 1 M imidazole. The eluate volume was reduced, and imidazole concentration lowered to ∼50 mM by sequential adding of buffer without imidazole using a 100 kDa cut-off concentrator (Amicon). NavMs proteins were treated with thrombin protease (Merck) for 16 h at 4°C then purified by size exclusion chromatography using a Superdex 200 10/300 column (Cytiva Life Sciences). NavMs was concentrated to 10 mg/ml and either used directly or stored at −80°C. NavMs mutants were concentrated to 10 mg/ml accounting for the different extinction coefficients caused by mutation.

### BTFA-labelling

Following purification, NavMs C52A/T19C protein was diluted to 10 μM. 5 mM BTFA (Merck) was added, and the mixture was incubated at 4°C for 24 h with gentle rotation. Free BTFA was removed from protein by gel filtration using a Superdex 200 10/300 column and gave a protein peak identical in elution volume when compared to untreated protein. Complete removal of the probe was confirmed by lack of a sharp peak at the BTFA fluorine resonance in the 1D ^19^F NMR spectrum of the NavMs-BTFA sample.

### Sevoflurane preparation

Sevoflurane concentrations used an aqueous concentration of 0.57 mM corresponding to twice the minimum alveolar concentration (2 MAC), where MAC (in vol%) is the concentration needed to prevent movement in 50% of subjects in response to a painful stimulus (comparable to the ED_50_).

### Structural techniques

#### Saturation Transfer Difference Nuclear Magnetic Resonance (STD-NMR)

^1^H Experiments were performed with 0.57 mM (2 MAC) sevoflurane in buffer (20 mM Tris-Cl, 300 mM NaCl, 0.52% Hega-10) +/− 5 μM NavMs protein and 5% D2O. ^19^F experiments we performed in the same conditions but using 50 μM fluorine-labelled NavMs protein. All STD NMR experiments were performed on a Bruker Biospin Avance IIIHD 700 spectrometer, equipped with a 5 mm TCI cryoprobe, using the acquisition software Topspin 3.5. NMR spectra were acquired by collecting alternating on- and off-resonance spectra with saturation achieved using a train of Gaussian-shaped pulses of 50 ms duration with a peak B1 field of 500 Hz determined over randomised collections at the saturation time point specified. Data were processed in Topspin 3.5 (Bruker) and fitted to the mono-exponential equation:

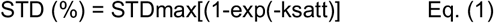

where STD (%) = (Voff – Von)/Voff x100. Voff and Von are the peak integrals from the off and on resonance saturation transfer NMR spectra. STDmax represents the STD at the maximal plateaued value.

^1^H STD NMR competition experiments were performed similarly to those described above but with 0.57 mM sevoflurane and 1 μM NavMs in mixtures containing competing concentrations of desflurane (∼20 mM), isoflurane (∼23 mM) or halothane (∼46 mM)

#### Crystallography

Crystallisation proceeded using stock NavMs F208L or mutants (10 mg/ml) in sitting drops at 4°C with drops containing 75 nl well condition and 75 nl NavMs dispensed using a mosquito® crystal robot (SPT Labtech). Apo-crystals were grown in well conditions containing 30-35% PEG 300,100 mM HEPES (pH 7) and 0-100 mM NaCl. For NavMs-SEVO crystals sevoflurane was saturated into reservoir solutions of crystal containing wells and incubated with the crystals for up to 1 h with periodic harvesting and flash freezing of individual crystals in liquid nitrogen.

#### Data Collection and Processing

Crystal optimisation was required in this study. X-ray diffraction data were collected for crystals using the beamlines P13 (EMBL, Hamburg, Germany), ID23-1 and ID30A-1 (ESRF, Grenoble, France) and I24 and I04 (Diamond Light Source, Oxford, UK) working at wavelengths between 0.886 and 0.9795 Å. Indexing and integration were performed with the XIA2-DIALS pipeline^61,62^, followed by scaling and merging with AIMLESS^63^. Molecular replacement was performed with PHASER^64^ using the search model (PDB ID: 6SX5) followed by model building in COOT^65^and structure refinement using REFMAC^66^. Polder Omit^44^ maps were created in Phenix^67^. Graphic illustrations were produced using PYMOL^68^ and UCSF Chimera^69^.

#### Long Wavelength X-ray Crystallography

Data were collected on the long-wavelength MX beamline I23 at Diamond Light Source at a wavelength of 2.755 Å (4.5 keV). This wavelength, although not optimal for chlorine (K-edge 4.4 Å, 2.82 keV), was selected as a compromise between the increasing signal from chlorine as it reaches its K-edge and the decreasing data quality that results from increasing X-ray absorption at longer wavelengths. Each dataset consisted of 360° rotation with an exposure time of 0.1 s per 0.1° image. Crystallographic data processing was performed as above. Anomalous difference maps were generated using the program Anode^70^.

### Electrophysiology

#### NavMs and NaChBac cDNA Constructs and Cell Culture

The NavMs-pTracer-CMV2 and NaChBac-pTracer-CMV3 plasmids carrying the separately expressed eGFP gene was used for whole-cell electrophysiology^33,71^. Standard site-directed mutagenesis protocols using the QuikChange Lightning kit (Qiagen, Germantown, MD) were used to create mutants in NavMs (Y143A and Y143F) and NaChBac (Y156A). The entire open reading frames of successful cDNA clones were confirmed by Sanger sequencing.

Mammalian HEK293T cells (CRL-3216 from ATCC, Manassas, VA) were maintained in high-glucose Dulbecco’s modified Eagle medium supplemented with 10% (vol/vol) fetal bovine serum (FBS), 2 mM GlutaMAX (Life Sciences, Waltham, MA) and 1% penicillin-streptomycin (Life Sciences). Passage numbers between 4 and 30 were used. Cells were seeded into 35-mm dishes and transfected with the respective NavMs cDNA after 1-3 days using Lipofectamine 2000 (Life Sciences) according to the manufacturer’s protocol. One day after transfection, cells were released with TrypsinLE (Life Sciences) and replated at lower density onto 12-mm round #1.5 glass coverslips (Warner Instruments, Holliston, MA) a minimum of 3 h before recording isolated adherent cells with eGFP fluorescence.

#### Human Nav1.1 cDNA Constructs and Cell Culture

Na^+^ currents from human neuronal Na_v_ subtype Na_v_1.1 were recorded and analysed by heterologous expression in a neuronal background using the ND7/23 neuroblastoma cell line^49^. An optimised version of WT human Na_v_1.1 (accession number NM_001165963) was kindly provided by A. L. George, Jr. (Northwestern University, Evanston, IL) and obtained via AddGene (Watertown, MA)^72^. Modification of hNav1.1 to be resistant to TTX (F383S) ^52^ allowed recordings of hNav1.1 currents in isolation of the endogenous Na^+^ current produced by these cells, which is completely blocked by 250 nM TTX. This TTX-resistant version has been shown to have no effect on gating properties, channel kinetics, or anaesthetic sensitivity^21,52^ and is referred to as hNa_v_1.1_R_.

Standard site-directed mutagenesis protocols using the QuikChange Lightning kit (Qiagen, Germantown, MD) were used to create the hNav1.1_R_ F902A and F1682A mutants. The entire open reading frames of successful cDNA clones were confirmed by Sanger sequencing.

Rodent ND7/23 neuroblastoma cells (Sigma-Aldrich, St. Louis, MO) were plated on 12-mm glass coverslips and incubated at 37°C in a humidified atmosphere of 5% CO_2_/95% air in Dulbecco’s modified Eagle’s medium supplemented with 10% (v/v) fetal bovine serum, 2 mM GlutaMax (Life Sciences), 1% penicillin-streptomycin (Life Sciences). This cell line provides a neuronal background for expression of Na_v_ by providing critical factors such as auxiliary subunits that facilitate functional expression of the channel protein^50^. ND7/23 cells were transfected with 4 μg hNa_v_1.1_R_ cDNA using Lipofectamine LTX (Invitrogen, Carlsbad, CA). Electrophysiological studies were conducted 24 h after transfection. TTX-resistant Na_v_ subtypes were studied in the presence of 250 nM TTX to block endogenous Na^+^ currents^21,50^.

#### Whole-cell Patch-clamp Electrophysiology

Pipettes were pulled from standard borosilicate glass (1.5 mm OD/0.86 mm ID; Sutter Instrument, Novato, CA) to a resistance of 3–4 MΩ (when filled) for NavMs or 1.5–2.5 MΩ for hNav1.1_R_ using a P-1000 puller (Sutter Instrument). Whole-cell voltage-clamp electrophysiology was performed^73^ using an AxoPatch 200B amplifier (Molecular Devices, San Jose, CA) connected to a DigiData 1550B analogue-to-digital converter (Molecular Devices). Signals were sampled at 20 kHz and filtered at 5 kHz, respectively. Series resistance was corrected 85–90%. Capacitive current transients were cancelled by the internal amplifier circuitry, and leak currents were subtracted using a standard P/4 protocol applied after the desired stimulus. For NavMs and NaChBac, cells were continuously perfused with extracellular solution at room temperature (24°C) containing (mM) 150 NaCl, 10 Hepes, 1.8 CaCl_2_ and 1 MgCl_2_, adjusted to pH 7.4 with NaOH. Pipette solutions contained (mM) 110 CsF, 30 NaCl, 10 Hepes, 5 ethylene glycol tetraacetic acid (EGTA), adjusted to pH 7.30 with CsOH. The osmolality of all solutions was balanced to 300±3 mOsm/kg H_2_O with sucrose. For hNav1.1_R_, cells were continuously perfused with extracellular solution at room temperature (24°C) containing (mM) 130 NaCl, 10 Hepes, 3.25 KCl, 2 CaCl_2_, 2 MgCl_2_, 0.1 CdCl_2_, 20 tetraethylammonium-Cl, 0.00025 tetrodotoxin, and 5 D-glucose adjusted to pH 7.4 with NaOH. Pipette solutions for hNav1.1_R_ contained (mM) 120 CsF, 10 NaCl, 10 Hepes, 10 ethylene glycol tetraaceticacid (EGTA), 10 tetraethylammonium-Cl, 1 CaCl_2_, 1 MgCl_2_ adjusted to pH 7.30 with CsOH. The osmolality of all solutions was balanced to 310±3 mOsm/kg H_2_O with sucrose. Saturated sevoflurane (Abbott Laboratories) stock solutions were prepared in extracellular solution in gas-tight glass vials and further diluted in gas-tight Hamilton syringes (Hamilton, Reno, NV) to the desired working concentration. Solutions in gas-tight syringes were delivered using a pressurised perfusion system (ALA Scientific Instruments, Farmingdale, NY) with Teflon tubing via a 200 μm diameter manifold tip positioned ∼200–300 μm from the recorded cell. After control recordings, sevoflurane was perfused before subsequent recordings and continuously thereafter until washout. Solutions were delivered through a pressurised perfusion system to minimise mechanical disturbance of cells during sevoflurane superfusion. Mock experiments with extracellular solution showed no effect on *I*_na_ or activation/inactivation kinetics (data not shown). Sevoflurane concentrations equivalent to clinically effective concentrations in mammals were 0.28 mM and 0.57 mM, equivalent to 1 or 2 times the minimum alveolar concentration (MAC) ^74^ after correction to room temperature and were confirmed by gas chromatography^50^.

### Molecular Docking

Molecular docking was performed using the docking platform Autodock Vina^75^ in UCSF Chimera^69^. Sevoflurane was docked into a box surrounding the sevoflurane binding site identified in NavMs (PDB ID: 6SX5) and the predicted cognate binding sites in human Nav1.1 (PDB ID: 7DTD) using a receptor search volume of 20×20×20 Å and selecting for generation of the top 10 binding modes for each search. Only the top binding modes produced at the sites in NavMs and Nav1.1 are reported here.

## Notes

https://doi.org/10.2210/pdb9gtq/pdb

https://doi.org/10.2210/pdb9gv1/pdb

### Competing Interest Statement

The authors have declared no competing interest.

### Summary of Updates

We have added new compelling data showing that mutation of a tyrosine residue in the homologous binding site in NaChBac completely removes the effect of sevoflurane on steady-state inactivation.

