## Supplementary Information for "Volatile anaesthetics modulate voltage-gated sodium channel function at a site critical for gating"

This PDF includes:

Figures S1 to S5
Tables S1 to S3

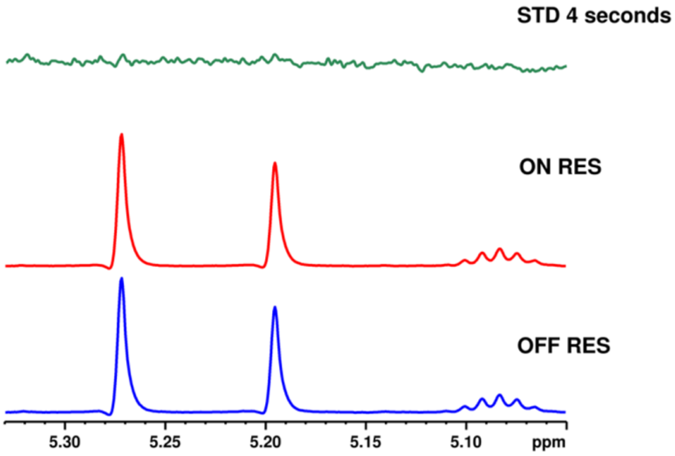

**Figure S1. STD at sevoflurane 1H resonances is not produced in the absence of NavMs.** Stacked spectra show that the sevoflurane 1H resonances are unaffected, and no STD is produced in STD NMR experiments performed with 2MAC sevoflurane in the absence of NavMs. The spectra are shown at the 4-second time point associated with STDmax in the same experiments performed with NavMs.

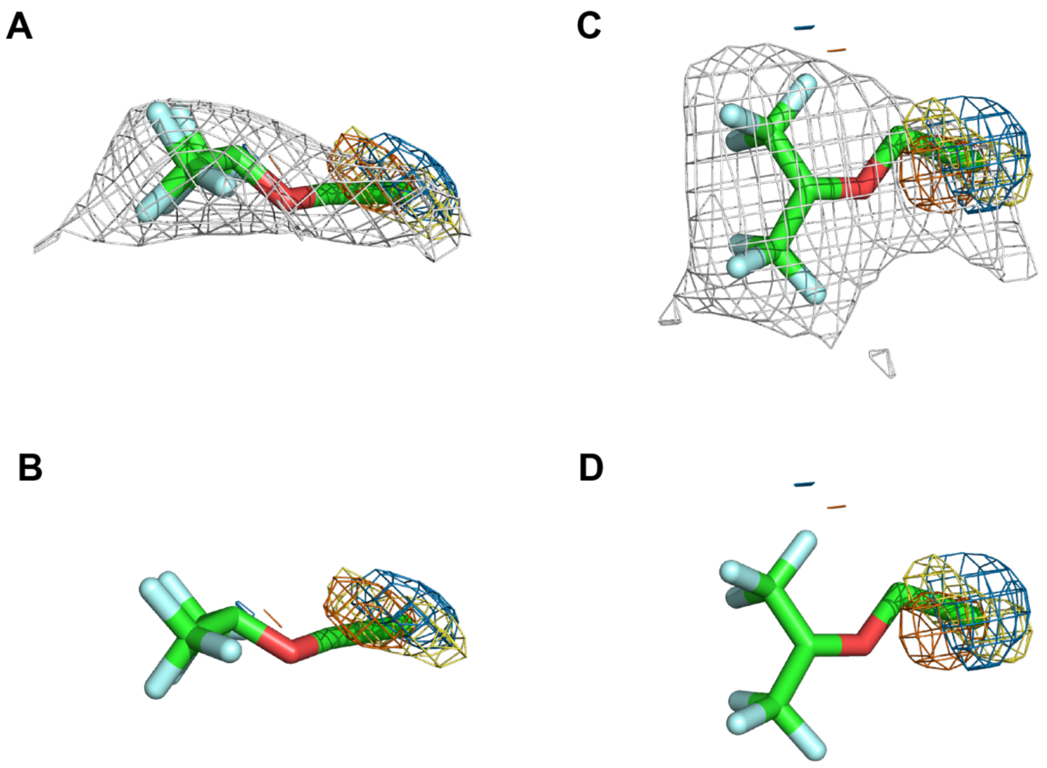

**Figure S2: A unique chlorine signal is found in the anomalous density maps from NavMs-chloroSEVO co-crystals.**
**a,** The unique signal identified in the anomalous maps of three NavMs-chloroSEVO co-crystals is represented as blue, red, and yellow mesh (contoured at 3σ). This signal is contained within the 2Fo-Fc electron density produced by chloroSEVO (shown as a grey mesh at 1σ), facilitating the accurate placement of chloroSEVO and sevoflurane into their density. **b,** Similar representation as in (**a**) but without the mesh for clarity. **c** and **d,** Alternative views of the densities presented in (**a**) and (**b**).

#### Supplementary Table S1: Activation and Inactivation of NavMs and NaChBac WT and mutant channels

| | | | | | | | |
|---|---|---|---|---|---|---|---|
| NavMs WT | *$V_{\frac{1}{2}Activ}$* | *Mean* | *SD* | *SEM* | *n* | *P value* | |
| 2 MAC | CTL | −71.5 | 7.6 | 2.7 | 8 | | |
| | SEVO | −66.7 | 6.6 | 2.3 | 8 | n.s. | 0.0695 |
| *Activation* | *Slope* | *Mean* | *SD* | *SEM* | *n* | *P value* | |
| | CTL | 11 | 2.2 | 0.8 | 8 | | n.s. |
| | SEVO | 13 | 2.8 | 1.0 | 8 | n.s. | 0.1606 |
| | *$V_{\frac{1}{2}Inact}$* | *Mean* | *SD* | *SEM* | *n* | *P value* | |
| | CTL | −106 | 6.4 | 2.2 | 8 | | |
| *Inactivation* | SEVO | −126 | 8.2 | 2.9 | 8 | **** | <0.0001 |
| | *Slope* | *Mean* | *SD* | *SEM* | *n* | *P value* | |
| | CTL | −8.4 | 1.6 | 0.6 | 8 | | |
| | SEVO | −11.9 | 1.9 | 0.7 | 8 | ** | 0.008 |
| NavMs | *$V_{\frac{1}{2}Activ}$* | *Mean* | *SD* | *SEM* | *n* | *P value* | |
| WT vs. Y143A | WT | −72.6 | 8.0 | 2.1 | 14 | | |
| | Y143A | −67.4 | 9.0 | 2.4 | 15 | n.s. | 0.1131 |
| *Activation* | *Slope* | *Mean* | *SD* | *SEM* | *n* | *P value* | |
| | WT | 11.5 | 2.7 | 0.7 | 14 | | |
| | Y143A | 14.6 | 3.1 | 0.8 | 15 | ** | 0.0079 |
| | *$V_{\frac{1}{2}Inact}$* | *Mean* | *SD* | *SEM* | *n* | *P value* | |
| | WT | −108 | 5.2 | 1.4 | 14 | | |
| *Inactivation* | Y143A | −129 | 9.5 | 2.2 | 18 | **** | <0.0001 |
| | *Slope* | *Mean* | *SD* | *SEM* | *n* | *P value* | |
| | WT | −10.08 | 2.9 | 0.8 | 14 | | |
| | Y143A | −16.85 | 4.6 | 1.1 | 18 | **** | <0.0001 |
| NavMs Y143F | *$V_{\frac{1}{2}Activ}$* | *Mean* | *SD* | *SEM* | *n* | *P value* | |
| 2 MAC | CTL | −76.7 | 9.0 | 4.0 | 5 | | |
| | SEVO | −79.6 | 9.7 | 4.4 | 5 | n.s. | 0.544 |
| *Activation* | *Slope* | *Mean* | *SD* | *SEM* | *n* | *P value* | |
| | CTL | 10.3 | 1.6 | 0.7 | 5 | | |
| | SEVO | 8.6 | 2.5 | 1.1 | 5 | n.s. | 0.391 |
| | *$V_{\frac{1}{2}Inact}$* | *Mean* | *SD* | *SEM* | *n* | *P value* | |
| | CTL | −107 | 7.9 | 3.6 | 5 | | |
| *Inactivation* | SEVO | −126 | 7.9 | 3.6 | 5 | * | 0.0187 |
| | *Slope* | *Mean* | *SD* | *SEM* | *n* | *P value* | |
| | CTL | −10.7 | 2.7 | 1.2 | 5 | | |
| | SEVO | −14.6 | 2.7 | 1.2 | 5 | * | 0.031 |

| | | | | | | | |
|---|---|---|---|---|---|---|---|
| NaChBac WT | $V_{½Activ}$ | *Mean* | *SD* | *SEM* | *n* | *P value* | |
| | CTL | −50.4 | 8.0 | 3.3 | 6 | | |
| *Activation* | SEVO | −53.2 | 9.3 | 3.8 | 6 | n.s. | 0.242 |
| *2 MAC* | *Slope* | *Mean* | *SD* | *SEM* | *n* | *P value* | |
| | CTL | 4.6 | 1.6 | 0.7 | 6 | | |
| | SEVO | 4.3 | 1.8 | 0.7 | 6 | n.s. | 0.703 |
| | $V_{½Inact}$ | *Mean* | *SD* | *SEM* | *n* | *P value* | |
| *Inactivation* | CTL | −60.8 | 4.0 | 1.5 | 7 | | |
| | SEVO | −74.8 | 6.1 | 2.3 | 7 | **** | <0.0001 |
| | *Slope* | *Mean* | *SD* | *SEM* | *n* | *P value* | |
| | CTL | −3.4 | 1.2 | 0.4 | 7 | | |
| | SEVO | −4.7 | 1.2 | 0.4 | 7 | n.s. | 0.090 |
| NaChBac Y156A | $V_{½Activ}$ | *Mean* | *SD* | *SEM* | *n* | *P value* | |
| | CTL | −73.6 | 8.2 | 2.7 | 9 | | |
| *Activation* | SEVO | −70.9 | 7.5 | 2.5 | 9 | n.s. | 0.279 |
| *2 MAC* | *Slope* | *Mean* | *SD* | *SEM* | *N* | *P value* | |
| | CTL | 13.0 | 1.4 | 0.5 | 5 | | |
| | SEVO | 11.8 | 1.9 | 0.6 | 5 | n.s. | 0.073 |
| | $V_{½Inact}$ | *Mean* | *SD* | *SEM* | *n* | *P value* | |
| *Inactivation* | CTL | −99.3 | 7.4 | 2.6 | 8 | | |
| | SEVO | −101 | 5.5 | 2.0 | 8 | n.s. | 0.140 |
| | *Slope* | *Mean* | *SD* | *SEM* | *n* | *P value* | |
| | CTL | −9.8 | 2.5 | 0.9 | 8 | | |
| | SEVO | −10.4 | 2.3 | 0.8 | 8 | n.s. | 0.428 |

#### Supplementary Table S2: Crystallographic Data and refinement Statistics

| ***Data collection*** | Apo-NavMs (F208L) PDB ID: 9GTQ | NavMs-SEVO (F208L) PDB ID: 9GV1 |
|---|---|---|
| Space group | I422 | I422 |
| *Unit-cell parameters* | | |
| a, b, c (Å) | 108.93,108.93, 209.37 | 108.70,108.70, 209.16 |
| α, β, γ (°) | 90, 90, 90 | 90, 90, 90 |
| Resolution (Å) | 2.20 | 2.2 |
| Completeness | 100.0 (100.0) | 100.0 (100.0) |
| *<I/σ(I)>* | 17.1 (3.3) | 28.6 (4.5) |
| CC(1/2) | 1.0 (0.954) | 0.999 (0.973) |
| $R_{merge\ all}$ | 0.099 | 0.069 |
| $R_{pim\ All}$ | 0.023 | 0.014 |
| Solvent content (%) | 61.1 | 59.0 |
| Molecule per ASU | 1 | 1 |
| Wilson B factor (Å$^2$) | 41.6 | 41.88 |
| ***Refinement*** | | |
| Resolution Range (Å) | 39.95 – 2.2 | 40.0 – 2.2 |
| $R_{work}$ | 0.215 | 0.225 |
| $R_{free}$ | 0.245 | 0.250 |
| Reflection, working | 32379 | 32200 |
| Reflection, free | 1665 | 1656 |
| Average B factor (all atoms) | 66.3 | 68.0 |
| Protein | 65.95 | 69.91 |
| Ligand/Ion | 86.39 | 88.9 |
| Water | 50.6 | 46.98 |
| RMS bond angle | 1.81 | 2.29 |
| RMS bond length (Å) | 0.0128 | 0.0127 |
| *Ramachandran Analysis:* | | |
| Preferred region (%) | 91.2 | 94.4 |
| Allowed region (%) | 8 | 5.2 |
| Outliers (%) | 0.8 | 0.4 |

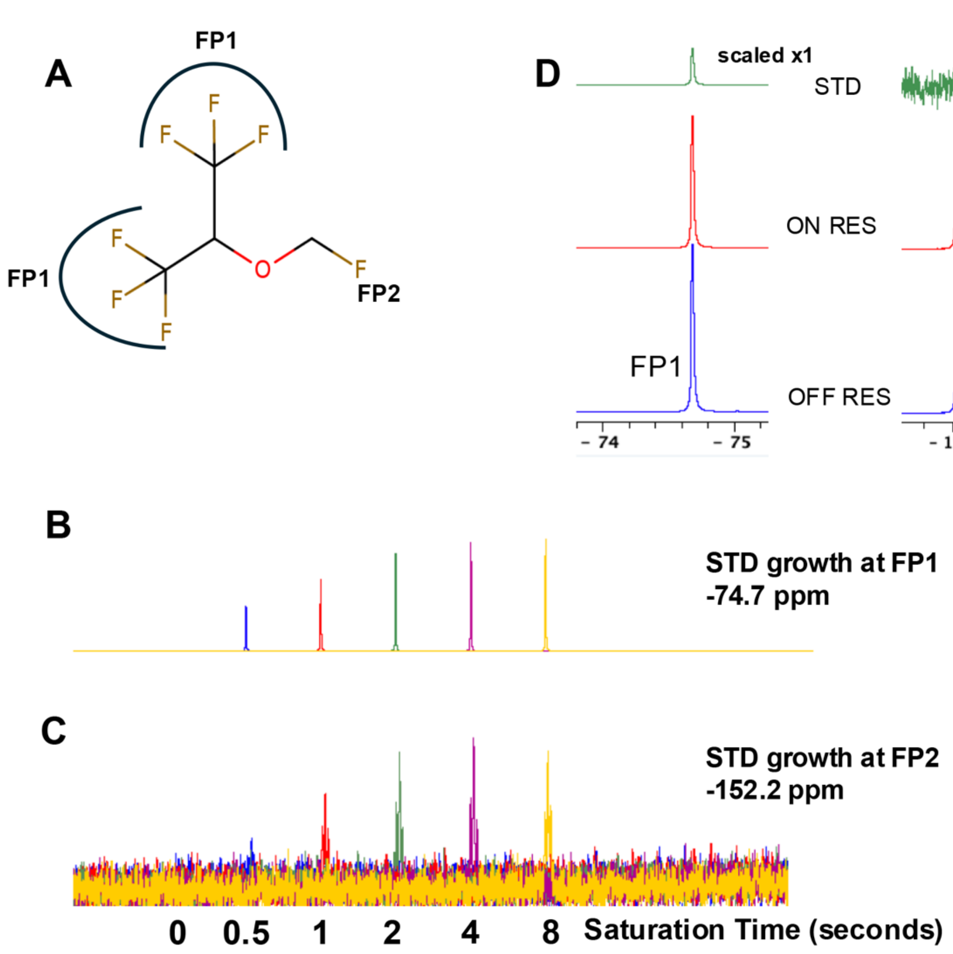

**Figure S3. $^{19}F$-$^{19}F$ Saturation Transfer from NavMs-T19C-BTFA to Sevoflurane.**
**a,** Structure of sevoflurane, showing the six equivalent trifluoromethyl fluorine atoms that produce a fluorine resonance peak (FP1) at −74.7 ppm and the single fluoromethyl fluorine that produces a fluorine resonance peak (FP2) at −152.2 ppm. **b,** STD build-up as a function of saturation time for the FP1 signal. **c,** STD build-up as a function of saturation time for the FP2 signal. **d,** Experiment at STDmax demonstrating that STD produced at the FP1 peak resonance (17.5%, *left*) is substantially greater to that produced at the FP2 peak resonance (3.5%, *right*). The FP2 peak is split into a triplet by J-coupling between the fluorine and hydrogen nuclei in the $-CH_2F$ group. STD (%) is calculated as (I_off − I_on) / I_off x 100, where I_off and I_on are the signal integrals of the sevoflurane fluorine peaks in the spectra at the OFF RES (0 ppm, empty of fluorine signal) and the ON RES (−83.8 ppm, which contains the unique fluorine signal from BTFA-labeled NavMs T19C) at indicated saturation times.

| | | | |
|---|---|---|---|
| NavMs | 237 | LLTVVFYIAAVMA | 249 |
| NaChBac | 250 | LMSIFFYIFAVIG | 262 |
| hNav1.1 DI | 253 | LTVFCLSVFALIG | 265 |
| hNav1.1 DII | 896 | VLAIIVFIFAVVG | 908 |
| hNav1.1 DIII | 1353 | VCLIFWLIFSIMG | 1365 |
| hNav1.1 DIV | 1676 | LLFLVMFIYAIFG | 1688 |

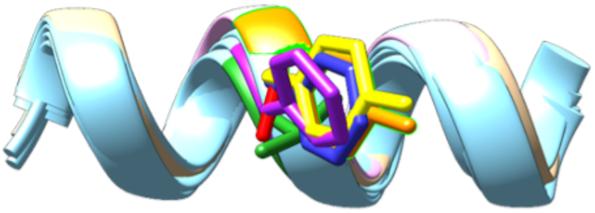

**Figure S4 Alignment of the S5 helix sequences relevant to this study.**
(*top*) Sections of S5 helices with colours highlighting Y143 from NavMs and the equivalent residues from NaChBac and each domain of human Nav1.1 (*bottom*) structural alignment of S5 helices with sidechains at the position of Y143 shown in colours matching alignment

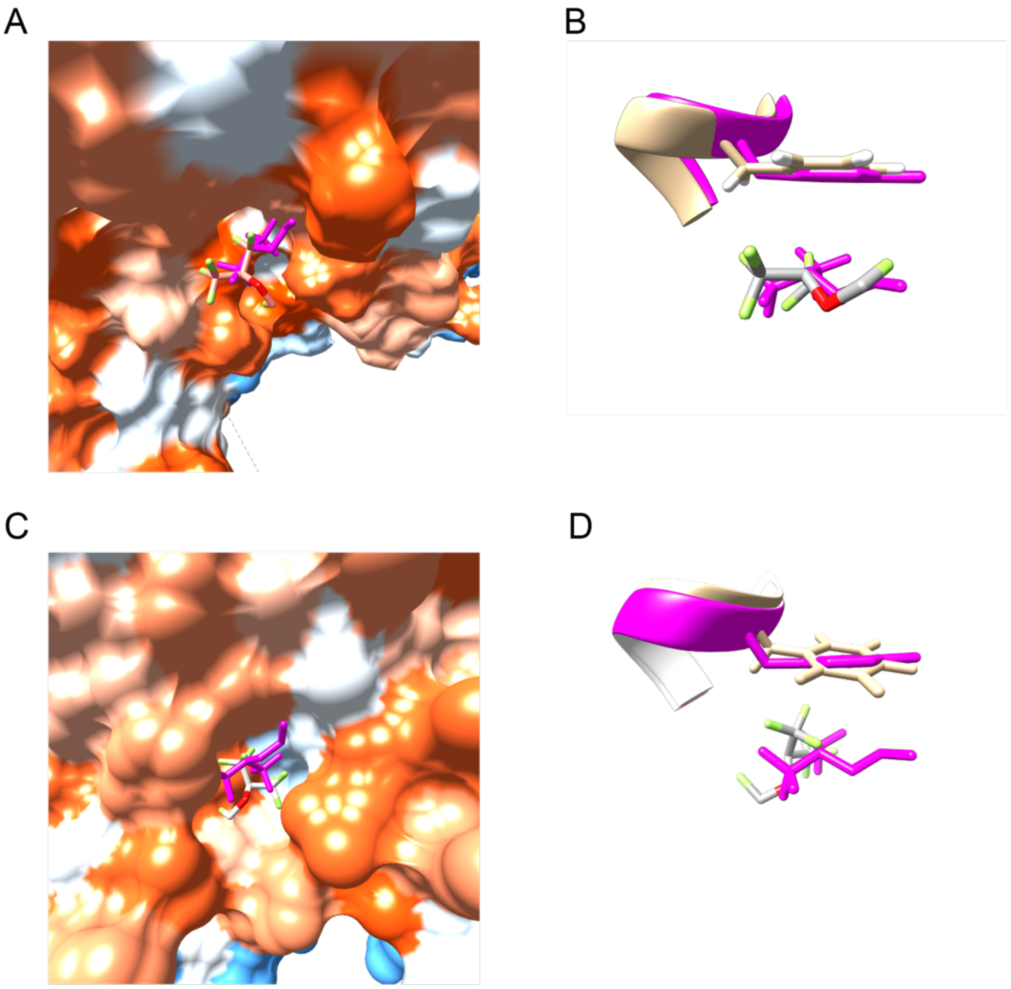

**Figure S5 Docking of sevoflurane into human Nav1.1 sites containing F902 and F1682.** Docking with AutoDock Vina generates sevoflurane poses that occupy pockets analogous to those found at the Y143 site of NavMs. **a,** Top binding pose for sevoflurane (stick representation, colored by heteroatom) in the DI-DII pocket of hNav1.1 containing F902. The relative position of sevoflurane (stick, magenta) in the NavMs crystal structure demonstrates an overlapping pose. **b,** Close up of (**a**), illustrating the positions of Y143 and sevoflurane in the crystal structure (magenta), overlaid F902 from human Nav1.1 and the top binding pose of sevoflurane from the simulation (colored by heteroatom). **c,** Top binding pose for sevoflurane (stick representation, colored by heteroatom) in the DIII-DIV pocket of human Nav1.1 containing F1682. The relative position of sevoflurane (stick, magenta) in the NavMs crystal structure demonstrates an overlapping pose. **d,** Close-up of (**c**), showing the positions of Y143 and sevoflurane in the crystal structure (magenta), overlaid with F1682 from human Nav1.1 and the top binding pose of sevoflurane from the simulation (colored by heteroatom).

### Table S3. Primer List for mutants used in this study

| Primers (Structural Biology) | Sequence (5' – 3') |
|---|---|
| C52A_For | GGAGCGTGTGGATCAACTTGCTCTGACTATCTTTATTGTTG |
| C52A_Rev | CAACAATAAAGATAGTCAGAGCAAGTTGATCCACACGCTCC |
| T19C_For | CGCTTTCAAAACGTCATCTGCGCCATTATTGTGCTC |
| T19C_Rev | GAGCACAATAATGGCGCAGATGACGTTTTGAAAGCG |
| Y143A_For | GTTGACGGTGGTCTTCGCGATTGCGGCTGTCATGG |
| Y143A_Rev | CCATGACAGCCGCAATCGCGAAGACCACCGTCAAC |
| Y143F_For | GTTGACGGTGGTCTTCTTCATTGCGGCTGTCATGG |
| Y143F_Rev | CCATGACAGCCGCAATGAAGAAGACCACCGTCAAC |
| Primers (NavMs Ephys) | Sequence (5' – 3') |
| Y143A_Fwd | CTGTTGACGGTGGTCTTCGCTATTGCGGCTGTCATGGC |
| Y143A_Rev | GCCATGACAGCCGCAATAGCGAAGACCACCGTCAACAG |
| Y143F_Fwd | GTTGACGGTGGTCTTCTTTATTGCGGCTGTCATGG |
| Y143F_Rev | CCATGACAGCCGCAATAAAGAAGACCACCGTCAAC |
| Primers (NaChBac Ephys) | Sequence (5' – 3') |
| Y156A_Fwd | CTTAATCTTGATGAGCATTTTCTTCGCTATTTTTGCCGTTATCGGGA CGATG |
| Y156A_Rev | CATCGTCCCGATAACGGCAAAAATAGCGAAGAAAATGCTCATCAA GATTAAG |
| Primers (hNav1.1 Ephys) | Sequence (5' – 3') |
| Nav1.1-F383S_Fwd | GTTTCGACTAATGACTCAGGACAGCTGGGAAAATCTTTATCAACTG |
| Nav1.1-F383S_Rev | CAGTTGATAAAGATTTTCCCAGCTGTCCTGAGTCATTAGTCGAAAC |
| Nav1.1R-F902A_Fwd | GTCTTGGCCATCATCGTCGCCATTTTTGCCGTGGTCGG |
| Nav1.1R-F902A_Rev | CCGACCACGGCAAAAATGGCGACGATGATGGCCAAGAC |
| Nav1.1R-F1682A_Fwd | CCTACTCTTCCTAGTCATGGCCATCTACGCCATCTTTGGG |
| Nav1.1R-F1682A_Rev | CCCAAAGATGGCGTAGATGGCCATGACTAGGAAGAGTAGG |
